# Evolutionary trajectories of β-lactam resistance in *Enterococcus faecalis* strains

**DOI:** 10.1101/2024.09.23.614543

**Authors:** Paul Ugalde Silva, Charlene Desbonnet, Louis B Rice, Mónica García-Solache

**Author notes:** Address correspondence to Mónica García-Solache.

## Abstract

Resistance to ampicillin and imipenem in *Enterococcus faecalis* is infrequent. However, the evolution of resistance can occur through prolonged antibiotic exposure during the treatment of chronic infections. In this study, we conducted a Long-Term Evolution Experiment (LTEE) using four genetically diverse strains of *E. faecalis* with varying susceptibilities to ampicillin and imipenem. Each strain was subjected to increasing concentrations of either ampicillin or imipenem over 200 days, with three independent replicates for each strain.

Selective pressure from imipenem led to the rapid selection of highly resistant lineages across all genetic backgrounds, compared to ampicillin. In addition to high resistance, we describe, for the first time, the evolution of a β-lactam dependent phenotype observed in lineages from all backgrounds. WGS and bioinformatic analysis revealed mutations in three main functional classes: genes involved in cell wall synthesis and degradation, genes in the walK/R two-component system, and genes in the c-di-AMP pathway. Our analysis identified new mutations in genes known to be involved in resistance as well as novel genes potentially associated with resistance.

Furthermore, the newly described β-lactam dependent phenotype was correlated with the inactivation of c-di-AMP degradation, resulting in high levels of this second messenger. Together, these data highlight the diverse genetic mechanisms underlying resistance to ampicillin and imipenem in *E. faecalis*. The emergence of high resistance levels and β-lactam dependency underscores the importance of understanding evolutionary dynamics in the development of antibiotic resistance.

**Importance:** *E. faecalis* is a major human pathogen, and treatment is frequently compromised by poor response to first-line antibiotics such ampicillin. Understanding the factors that play a role in susceptibility/resistance to these drugs will help guide the development of much needed treatments.

## Introduction

*Enterococcus faecalis* is one of the leading causes of healthcare-associated infections (1, 2). The widespread use of antibiotics in various human activities has led to the selection of strains equipped with mechanisms to resist antibiotic treatments to which they were previously susceptible. Increasing emergence of antibiotic resistance in this commonly occurring pathogen (3) represents a clinical challenge and is associated with worse clinical outcomes when compared with susceptible strains (4, 5). Mortality rates of enterococcal bacteremia range from 20% to 50%, and patient comorbidities and antimicrobial resistance are the major risk factors for early mortality (6).

*E. faecalis* expresses inherent resistance to many β-lactam treatments, although the extent of resistance varies depending on the specific class of β-lactam. Amino-penicillins, such as ampicillin, are highly effective, while carbapenems, like imipenem, exhibit lower efficacy, and cephalosporins, are the least effective (7). In clinical practice, ampicillin is considered the treatment of choice for infections caused by susceptible *E. faecalis* strains (8), however, we and others have reported clinical strains showing ampicillin resistance (9–12). Imipenem, although not specifically indicated for enterococcal infections, demonstrates in vitro activity against many strains and so is sometimes used (13–15).

Development of resistance to ampicillin and imipenem during typical courses of antimicrobial therapy is rare in *E. faecalis* (16, 17). However, the evolution of resistance to β-lactams may be associated with prolonged antibiotic exposure during the treatment of chronic infections. Previously, we reported a clinical isolate of *E. faecalis* obtained from a patient who had undergone extended amoxicillin therapy, that was resistant to ampicillin and penicillin. The isolate carried mutations in one of the main determinants for β-lactam resistance, the penicillin binding protein 4 gene (*pbp4*). These changes led to a reduction in Pbp4’s affinity for β-lactams and triggered its overproduction (9). While mutations in *pbp4* conferred an advantage during selection pressure resulting in emergence of resistance, they were associated with reduced bacterial growth rate and cell wall stability.

Resistance and the associated fitness costs may arise from diverse evolutionary trajectories, influenced by the cooperative effects of acquired mutations and the underlying genetic backgrounds (18–20). Consequently, predicting the specific trajectories leading to the evolution of resistance poses a significant challenge (21, 22). Aside from *pbp4* mutations, little is known about additional mechanisms of ampicillin resistance in *E. faecalis*. Our goal in the present study was to identify convergent and novel pathways to β-lactam resistance in *E. faecalis* strains with diverse genetic backgrounds.

To understand the evolutionary trajectories leading to β-lactam resistance in *E. faecalis*, we conducted a long-term evolution experiment (LTEE) using four distinct strains originating from different sources subjected to escalating concentrations of either ampicillin or imipenem over a period of 200 days. Through this experiment, we validated previously described genes leading to resistance to β-lactams, identified novel pathways and determined that although there were convergent pathways each strain follow a different evolutionary trajectory. Our findings underscore the significance of genetic background in predicting specific resistance patterns and highlight the genetic changes associated with fitness costs imposed during the evolution of resistance.

## Results

### Selection of β-lactam resistant *E. faecalis* in a LTEE

We set up a LTEE using four different *E. faecalis* strains from different sources and genetic backgrounds (Table 1). We obtained a total of 432 samples from our initial passages at the end of the experiment. Additionally, we inoculated a separate plate to initiate a parallel LTEE (replica cultures) on day 68 to investigate the reproducibility of the path to resistance in each strain. From the replicate cultures we obtained a total of 252 samples. The total number of antibiotic-selected lineages obtained in our experiment was 48 (Table S1).

**Table 1.** Strain information.

| **Strain** | **Source** | **Amp MIC** | **Imi MIC** | **Reference** |
| --- | --- | --- | --- | --- |
| **LS4828** | human - prosthetic knee joint | 16 | 16 | 9 |
| **ATCC 29212** | human - urine | 2 | 2 | 23 |
| **JH2-2** | human - stools | 1.5 | 1.5 | 24 |
| **D32** | porcine - stools | 2 | 2 | 25 |

### Evolution of β-lactam resistance during the LTEE

Periodic ampicillin and imipenem susceptibility testing during the experiment revealed a progressive loss of susceptibility to ampicillin and imipenem under both antibiotics’ selection.

Ampicillin exposure selected for ampicillin resistance in the three ATCC 29212 and D32 lineages and in two of three JH2-2 lineages in both original and replica plates, additionally all ampicillin-selected lineages evolved imipenem resistance (Table 2). Resistance to ampicillin and imipenem (MIC ≥ 8 µg/ml) in the lineages from the susceptible strains ATCC 29212 (23), JH2-2 (24), and D32 (25) appeared later during the experiment (around passage 122).

**Table 2.**
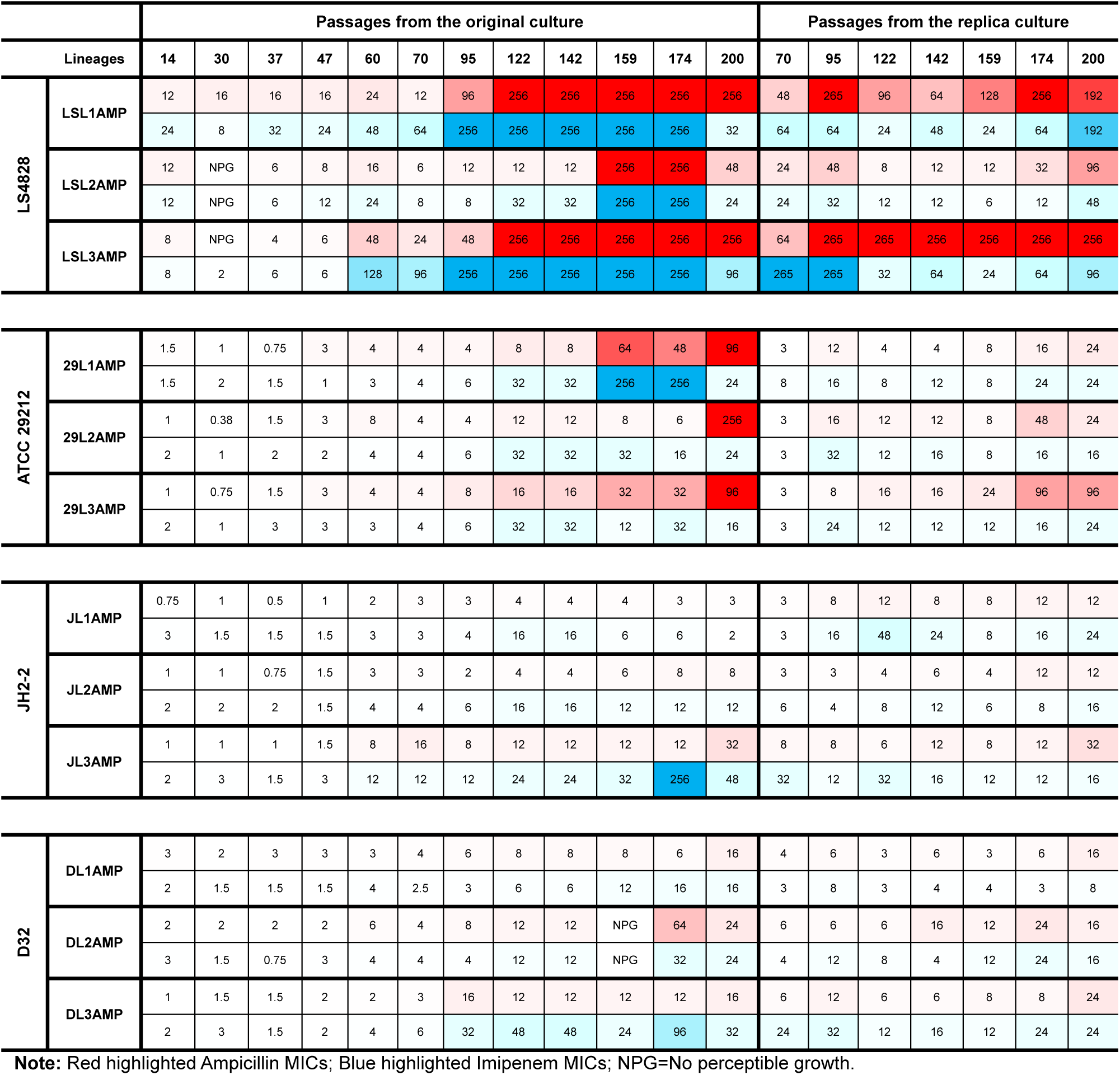
Ampicillin and imipenem MICs from lineages selected with ampicillin. Heat map of the Minimal Inhibitory Concentration (MIC) of ampicillin (red) and imipenem (blue) obtained from the lineages evolved under ampicillin exposure. MICs were determined between 18-24h of incubation and read by eye. The vertical dotted line delimits the passages from the original culture and the passages from the replica culture. Squares with a diagonal solid line reflect passages at which the MIC could not be obtained between 18-24h. Strain designation: LS lineages are derived from LS4828; 29 lineages are derived from ATCC 29212, J lineages are derived from JH2-2; D lineages are derived from D32. Lineage designation: First initial describes strain name, second initial and number describes lineage number; amp or imi informs antibiotic selection.

|  |  | Passages from the original culture |  |  |  |  |  |  |  |  |  |  |  | Passages from the replica culture |  |  |  |  |  |  |
| --- | --- | --- | --- | --- | --- | --- | --- | --- | --- | --- | --- | --- | --- | --- | --- | --- | --- | --- | --- | --- |
| Lineages |  | 14 | 30 | 37 | 47 | 60 | 70 | 95 | 122 | 142 | 159 | 174 | 200 | 70 | 95 | 122 | 142 | 159 | 174 | 200 |
| LS4828 | LSL1AMP | 12 | 16 | 16 | 16 | 24 | 12 | 96 | 256 | 256 | 256 | 256 | 256 | 48 | 265 | 96 | 64 | 128 | 256 | 192 |
|  |  | 24 | 8 | 32 | 24 | 48 | 64 | 256 | 256 | 256 | 256 | 256 | 32 | 64 | 64 | 24 | 48 | 24 | 64 | 192 |
|  | LSL2AMP | 12 | NPG | 6 | 8 | 16 | 6 | 12 | 12 | 12 | 256 | 256 | 48 | 24 | 48 | 8 | 12 | 12 | 32 | 96 |
|  |  | 12 | NPG | 6 | 12 | 24 | 8 | 8 | 32 | 32 | 256 | 256 | 24 | 24 | 32 | 12 | 12 | 6 | 12 | 48 |
|  | LSL3AMP | 8 | NPG | 4 | 6 | 48 | 24 | 48 | 256 | 256 | 256 | 256 | 256 | 64 | 265 | 265 | 256 | 256 | 256 | 256 |
|  |  | 8 | 2 | 6 | 6 | 128 | 96 | 256 | 256 | 256 | 256 | 256 | 96 | 265 | 265 | 32 | 64 | 24 | 64 | 96 |
| ATCC 29212 | 29L1AMP | 1.5 | 1 | 0.75 | 3 | 4 | 4 | 4 | 8 | 8 | 64 | 48 | 96 | 3 | 12 | 4 | 4 | 8 | 16 | 24 |
|  |  | 1.5 | 2 | 1.5 | 1 | 3 | 4 | 6 | 32 | 32 | 256 | 256 | 24 | 8 | 16 | 8 | 12 | 8 | 24 | 24 |
|  | 29L2AMP | 1 | 0.38 | 1.5 | 3 | 8 | 4 | 4 | 12 | 12 | 8 | 6 | 256 | 3 | 16 | 12 | 12 | 8 | 48 | 24 |
|  |  | 2 | 1 | 2 | 2 | 4 | 4 | 6 | 32 | 32 | 32 | 16 | 24 | 3 | 32 | 12 | 16 | 8 | 16 | 16 |
|  | 29L3AMP | 1 | 0.75 | 1.5 | 3 | 4 | 4 | 8 | 16 | 16 | 32 | 32 | 96 | 3 | 8 | 16 | 16 | 24 | 96 | 96 |
|  |  | 2 | 1 | 3 | 3 | 3 | 4 | 6 | 32 | 32 | 12 | 32 | 16 | 3 | 24 | 12 | 12 | 12 | 16 | 24 |
| JH2-2 | JL1AMP | 0.75 | 1 | 0.5 | 1 | 2 | 3 | 3 | 4 | 4 | 4 | 3 | 3 | 3 | 8 | 12 | 8 | 8 | 12 | 12 |
|  |  | 3 | 1.5 | 1.5 | 1.5 | 3 | 3 | 4 | 16 | 16 | 6 | 6 | 2 | 3 | 16 | 48 | 24 | 8 | 16 | 24 |
|  | JL2AMP | 1 | 1 | 0.75 | 1.5 | 3 | 3 | 2 | 4 | 4 | 6 | 8 | 8 | 3 | 3 | 4 | 6 | 4 | 12 | 12 |
|  |  | 2 | 2 | 2 | 1.5 | 4 | 4 | 6 | 16 | 16 | 12 | 12 | 12 | 6 | 4 | 8 | 12 | 6 | 8 | 16 |
|  | JL3AMP | 1 | 1 | 1 | 1.5 | 8 | 16 | 8 | 12 | 12 | 12 | 12 | 32 | 8 | 8 | 6 | 12 | 8 | 12 | 32 |
|  |  | 2 | 3 | 1.5 | 3 | 12 | 12 | 12 | 24 | 24 | 32 | 256 | 48 | 32 | 12 | 32 | 16 | 12 | 12 | 16 |
| D32 | DL1AMP | 3 | 2 | 3 | 3 | 3 | 4 | 6 | 8 | 8 | 8 | 6 | 16 | 4 | 6 | 3 | 6 | 3 | 6 | 16 |
|  |  | 2 | 1.5 | 1.5 | 1.5 | 4 | 2.5 | 3 | 6 | 6 | 12 | 16 | 16 | 3 | 8 | 3 | 4 | 4 | 3 | 8 |
|  | DL2AMP | 2 | 2 | 2 | 2 | 6 | 4 | 8 | 12 | 12 | NPG | 64 | 24 | 6 | 6 | 6 | 16 | 12 | 24 | 16 |
|  |  | 3 | 1.5 | 0.75 | 3 | 4 | 4 | 4 | 12 | 12 | NPG | 32 | 24 | 4 | 12 | 8 | 4 | 12 | 24 | 16 |
|  | DL3AMP | 1 | 1.5 | 1.5 | 2 | 2 | 3 | 16 | 12 | 12 | 12 | 12 | 16 | 6 | 12 | 6 | 6 | 8 | 8 | 24 |
|  |  | 2 | 3 | 1.5 | 2 | 4 | 6 | 32 | 48 | 48 | 24 | 96 | 32 | 24 | 32 | 12 | 16 | 12 | 24 | 24 |

**Note:** Red highlighted Ampicillin MICs; Blue highlighted Imipenem MICs; NPG=No perceptible growth.

Lineages under imipenem selection acquired higher MICs to both imipenem and ampicillin earlier in the experiment compared to ampicillin selection (Table 3). In the susceptible backgrounds, most displayed higher MICs to imipenem than to ampicillin in both original and replica cultures. Cross-resistance to ampicillin evolved in most lineages (Table 3); resistance to ampicillin in D32-derived lineages was slow to evolve compared to the other backgrounds. Interestingly, the break point (≥ 8 µg/ml) was reached in most cases between passages 95 and 122.

**Table 3.** Ampicillin and imipenem MICs from lineages selected with imipenem. Heat map of the Minimal Inhibitory Concentration (MIC) of ampicillin (red) and imipenem (blue) obtained from the lineages evolved under imipenem exposure. MICs were determined between 18-24h of incubation and read by eye. The vertical dotted line delimits the passages from the original culture and the passages from the replica culture. Squares with a diagonal solid line reflect passages which the MIC could not be obtained between 18-24h.

|  |  | Passages from the original culture |  |  |  |  |  |  |  |  |  |  |  | Passages from the replica culture |  |  |  |  |  |  |
| --- | --- | --- | --- | --- | --- | --- | --- | --- | --- | --- | --- | --- | --- | --- | --- | --- | --- | --- | --- | --- |
| Lineages |  | 14 | 30 | 37 | 47 | 60 | 70 | 95 | 122 | 142 | 159 | 174 | 200 | 70 | 95 | 122 | 142 | 159 | 174 | 200 |
| LS4828 | LSL1IMI | 16 | NPG | 6 | 24 | 32 | 32 | 256 | 256 | 256 | 256 | 256 | 256 | 48 | 265 | 96 | 48 | 256 | 256 | 256 |
|  |  | 24 | NPG | 32 | 24 | 96 | 192 | 256 | 256 | 256 | 256 | 256 | 256 | 265 | 265 | 96 | 64 | 256 | 256 | 256 |
|  | LSL2IMI | 16 | 3 | 8 | 16 | 24 | 32 | 256 | 256 | 256 | 256 | 256 | 256 | 48 | 265 | 48 | 64 | 256 | 257 | 256 |
|  |  | 12 | 8 | 8 | 32 | 256 | 256 | 256 | 256 | 256 | 256 | 256 | 256 | 265 | 265 | 128 | 256 | 48 | 64 | 256 |
|  | LSL3IMI | 16 | 3 | 6 | 24 | 16 | 32 | 256 | 256 | 256 | 256 | 256 | 256 | 64 | 265 | 265 | 256 | 256 | 256 | 256 |
|  |  | 24 | 12 | 24 | 32 | 32 | 64 | 256 | 256 | 256 | 256 | 256 | 256 | 265 | 265 | 265 | 48 | 48 | 192 | 256 |
| ATCC 29212 | 29L1IMI | 0.38 | 0.5 | 0.75 | 4 | 8 | 4 | 16 | 256 | 256 | 256 | 256 | 256 | 8 | 24 | 32 | 24 | 8 | 32 | 12 |
|  |  | 1.5 | 1 | 2 | 4 | 6 | 8 | 32 | 256 | 256 | 256 | 256 | 32 | 16 | 48 | 32 | 48 | 16 | 48 | 256 |
|  | 29L2IMI | 0.5 | 0.38 | 0.75 | 1.5 | 3 | 4 | 12 | 16 | 16 | 32 | 6 | 24 | 3 | 4 | 8 | 12 | 8 | 12 | 24 |
|  |  | 1.5 | 1 | 3 | 3 | 2 | 6 | 32 | 256 | 256 | 256 | 256 | 64 | 12 | 16 | 16 | 16 | 12 | 48 | 48 |
|  | 29L3IMI | 0.75 | 0.5 | 1 | 1.5 | 4 | 6 | 4 | 4 | 4 | 24 | 16 | 32 | 4 | 8 | 6 | 4 | 3 | 16 | 12 |
|  |  | 2 | 1.5 | 3 | 3 | 4 | 32 | 16 | 48 | 48 | 96 | 256 | 48 | 12 | 24 | 16 | 16 | 16 | 32 | 48 |
| JH2-2 | JL1IMI | 1.5 | 1.5 | 2 | 3 | 8 | 4 | NPG | NPG | NPG | NPG | NPG | NPG | 8 | 24 | 16 | 96 | 48 | 256 | 48 |
|  |  | 2 | 4 | 3 | 2 | 4 | 6 | NPG | NPG | NPG | NPG | 256 | 256 | 12 | 32 | 96 | 24 | 16 | 48 | 48 |
|  | JL2IMI | 0.75 | 1 | 1.5 | 2 | 2 | 8 | 12 | 12 | 12 | 48 | 64 | 256 | 6 | 6 | 12 | 8 | 6 | 8 | 12 |
|  |  | 2 | 3 | 4 | 2 | 3 | 32 | 16 | 32 | 32 | 64 | 256 | 256 | 16 | 16 | 64 | 24 | 24 | 32 | 32 |
|  | JL3IMI | 1.5 | 0.5 | 1 | 4 | 8 | 16 | 12 | 12 | 12 | 16 | 8 | 16 | 8 | 6 | 8 | 16 | 6 | 12 | 8 |
|  |  | 3 | 1.5 | 1.5 | 4 | 32 | 32 | 64 | 256 | 256 | 256 | 256 | 256 | 24 | 64 | 256 | 48 | 32 | 64 | 256 |
| D32 | DL1IMI | 1.5 | 1 | 0.75 | 4 | 6 | 7 | 6 | 6 | 6 | 4 | 4 | 8 | 3 | 6 | 4 | 6 | 6 | 6 | 8 |
|  |  | 3 | 2 | 2 | 3 | 8 | 32 | 64 | 256 | 256 | 256 | 256 | 256 | 16 | 32 | 24 | 24 | 32 | 48 | 48 |
|  | DL2IMI | 1 | 1 | 1 | 2 | 2 | 3 | 4 | 4 | 4 | 4 | 6 | 8 | 2 | 32 | 6 | 3 | 3 | 4 | 4 |
|  |  | 2 | 3 | 3 | 3 | 3 | 6 | 32 | 32 | 32 | 32 | 32 | 32 | 6 | 12 | 12 | 12 | 8 | 12 | 48 |
|  | DL3IMI | 1 | 2 | 1.5 | 6 | 3 | 10 | 12 | 8 | 8 | 16 | 12 | 24 | 8 | 12 | 12 | 12 | 12 | 12 | 16 |
|  |  | 1.5 | 4 | 1.5 | 4 | 8 | 32 | 48 | 256 | 256 | 256 | 256 | 256 | 32 | 48 | 32 | 64 | 32 | 48 | 64 |

**Note:** Red highlighted Ampicillin MICs; Blue highlighted Imipenem MICs; NPG=No perceptible growth.

LS4828, which was resistant to ampicillin and imipenem at the start, evolved increased resistance to both antibiotics, reaching MICs ≥ 256 µg/ml.

Our data indicate that continuous imipenem exposure selects for the evolution of high-level ampicillin and imipenem resistance at higher frequency than ampicillin exposure. Imipenem-selected lineages show higher variability in the MIC distribution and timing of β-lactam resistance acquisition. Imipenem selection tends to favor the rapid evolution of highly resistant lineages, compared to ampicillin selective pressure, across all tested genetic backgrounds. Evolution of ampicillin resistance seems to carry stronger fitness costs than imipenem exposure, since it is slower to evolve, and the MICs tend to be lower than lineages evolved under imipenem.

In the lineages passaged under no antibiotic pressure (Table S2), two lineages from the resistant strain LS4828 lost resistance to ampicillin and imipenem (MICs between 0.5 to 3 µg/ml). The ampicillin and imipenem MICs of the lineages from the susceptible backgrounds did not change.

Our findings demonstrate a remarkable ability of *E. faecalis* to adapt to elevated concentrations of β-lactams while also losing resistance if the selection pressure is removed, suggesting that β-lactam resistance carries fitness costs, as demonstrated by a lower growth rate of LS4828 and all resistant lineages compared to susceptible strains (data not shown).

### Chronic exposure to ampicillin and imipenem selected the evolution of β-lactam dependence

During the LTEE, in addition to the evolution of β-lactam resistance, we identified an unexpected β-lactam-dependent phenotype in 30 resistant lineages from all genetic backgrounds. This phenotype was observed as a dense zone of bacterial growth (growth halo) near the highest antibiotic concentrations of the MIC test strip, with little to no colonies observed in the proximity of the lower antibiotic concentrations or away from the strip (Fig. 1A). Dependence on the antibiotic to grow was observed as early as passage 70 and after the acquisition of resistance. In 7 of the lineages, it was later lost (Fig. 1B). In general, the dependent phenotype was more stable in later passages, with less colonies appearing on the periphery of the plate away from the disc (data not shown).

**Figure 1.**
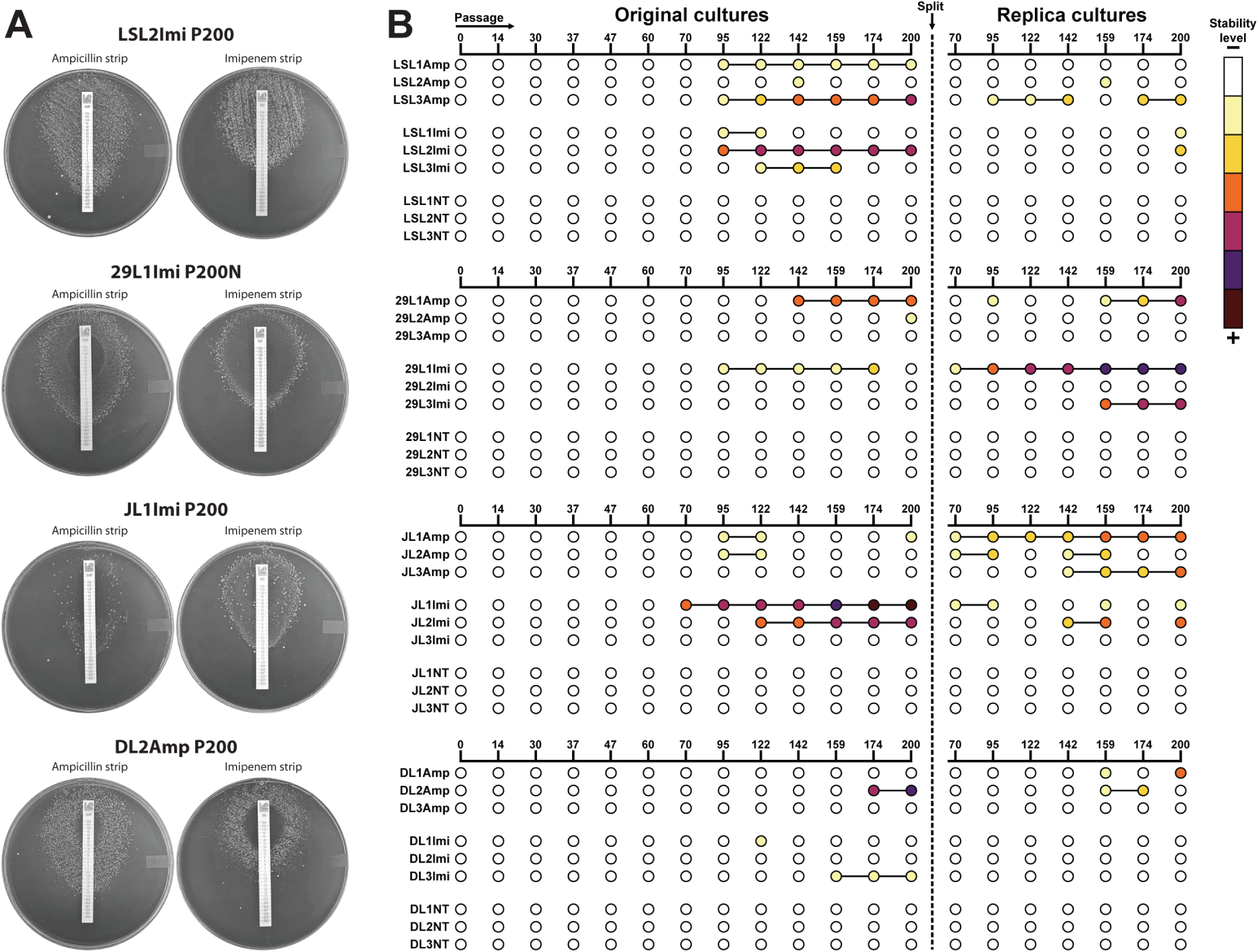
β-lactam dependent lineages. **A)** MIC strip tests of strongly dependent passage 200 lineages. LSL2Imi P200 (LS4828), 29L1Imi P200R (ATCC29212, replica culture), JL1Imi P200 (JH2-2), and DL2Amp P200 (D32). The plates show dense areas of growth near the test strip and an area devoid of colonies in the lower antibiotic concentrations and away from the strip. **B)** Dot chart highlighting β-lactam dependence during the experiment. Light to dark color shows the progression of the phenotype and how stable it was as determined by: 1) dependence is present at 24 h, but lost at 72h and all colonies are homogenous; 2) dependence is present at 24 h, but lost at 72 h, two types of colonies; 3) dependence is present at 24 and 72h, two types of colonies.

To test the stability of the dependent phenotype we performed growth curves with CFU counts for 4 antibiotic-dependent lineages at passage 200, cultures were supplemented with either ampicillin or imipenem, or grown without antibiotic (Fig. S2A). The dependent lineages grew faster in the presence of either ampicillin or imipenem, although ampicillin-grown cells presented a slower growth rate and reached lower CFUs (Fig. S2A). The absence of antibiotics negatively impacted the growth of the four lineages. The lineages grown in the presence of β-lactams maintained a dependent phenotype, forming growth halos around the discs (ampicillin, imipenem and penicillin). The four lineages lost antibiotic dependence when they were grown in the absence of β-lactams. All lineages maintained β-lactam resistance (Fig. S2B).

Since selection with ampicillin or imipenem resulted in cross resistance to the other antibiotic and to penicillin, we tested by disc diffusion assay thirteen different β-lactams for cross-resistance and dependence against four dependent lineages (table S3). The four lineages formed robust growth halos around penicillin, ampicillin, and imipenem discs, and around two other penicillins: ticarcillin and piperacillin. Growth around cephalosporins was more variable, less robust and slower. None of the lineages grew around oxacillin, cefotetan, or vancomycin discs (Table S3).

### Genome evolution under ampicillin and imipenem selection

We sequenced a total of 135 genomes: 46 derived from LS4828, 21 from D32, 24 from ATCC 29212 and 44 from JH2-2 from our LTEE, but we discarded four that were found to be a cross-contamination (Table S1). We also sequenced our laboratory stocks for the parental strains, for accurate variant calling.

Lineages derived from the resistant strain LS4828, accumulated less mutations compared with the lineages derived from the susceptible strains (Fig. S3). Deletions >50 bp were rare in most genomes, except for DL2Amp which lost a 12,895 bp plasmid, losing 16 genes. After culture split, DL2AmpR started to accumulate mutations between passage 95 and 122, coincidental with a 10 bp deletion in the DNA mismatch repair protein MutS gene. Mutations in *mutS* and *mutL* have been correlated with an increase in the spontaneous mutation rate in *E. faecalis* (26). Although each susceptible strain followed a different evolutionary trajectory to acquire β-lactam resistance we identified a core of genes mutated in more than one lineage (Fig. 2), suggesting their importance in either resistance or fitness as described below.

**Figure 2.**
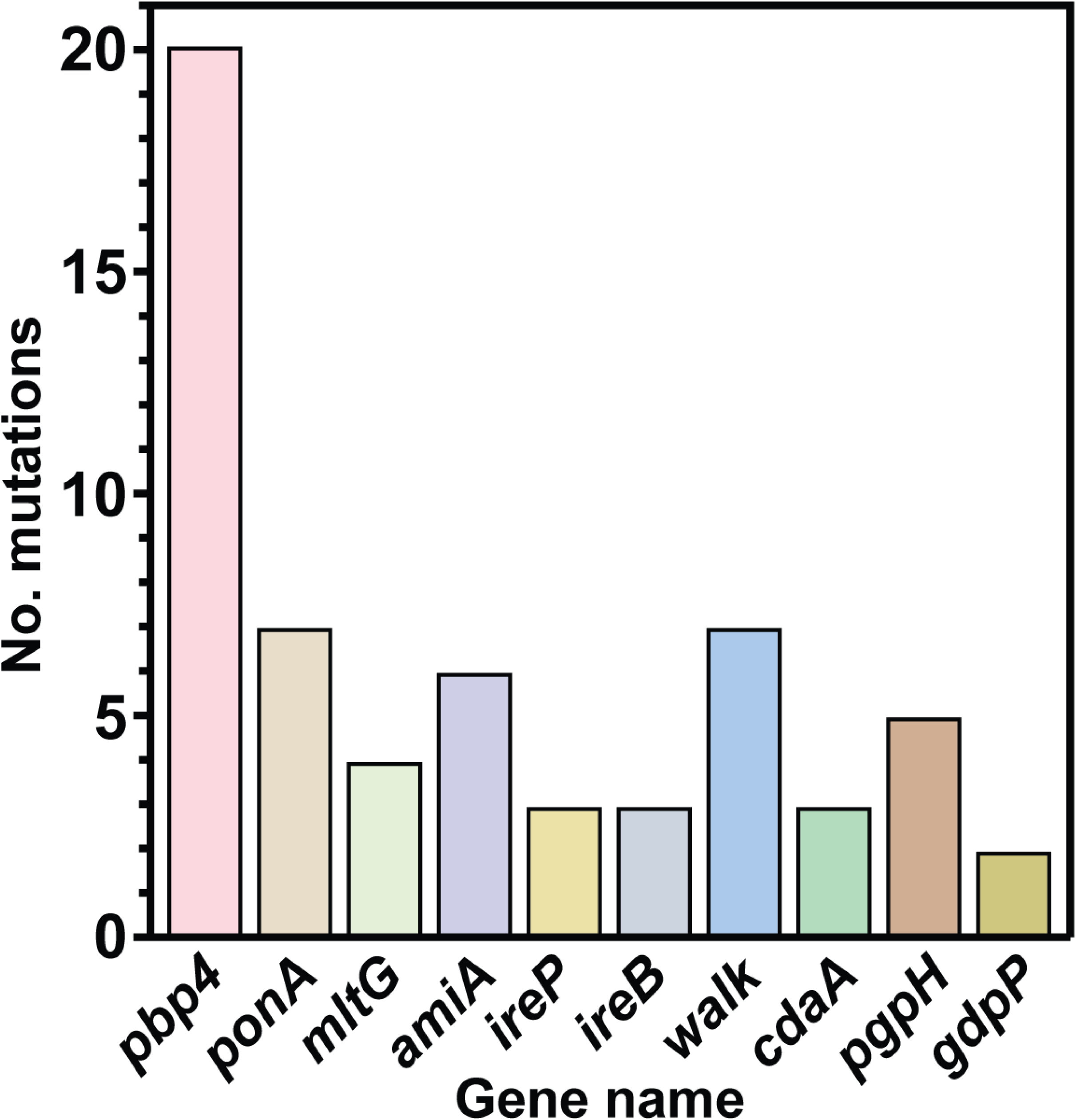
Mutations in cell wall regulating genes. The graph shows the number of mutations identified in genes involved in cell regulation that were present in more than one lineage.

Our LTEE allowed us to describe for the first time the mutational steps involved in the evolution of very high levels of resistance to ampicillin and imipenem (MICs ≥ 50µg/ml) in *E. faecalis*.

### Mutations in genes related to cell wall metabolism

The main target for β-lactams are the cell wall-building penicillin binding proteins (Pbp’s). During the experimental evolution of β-lactam resistance, we identified different mutations affecting the genes for these enzymes and other genes involved in cell wall homeostasis in all the sequenced lineages (Fig. 2, Table S4).

In *E. faecalis,* Pbp4 has been described as the main determinant for β-lactam resistance, and as such we identified mutations in the *pbp4* gene in resistant lineages derived from the three susceptible backgrounds, we did not identify new *pbp4* mutations in lineages derived from LS4828. We found two different kinds of mutations: those affecting the promoter region and those affecting the coding region. We identified three mutation sites in the promoter: the deletion of an A at positions - 57/-51 and -88/-81, and a nucleotide substitution at position -46 (relative to the start codon of the gene) (Fig. S4A).

In *pbp4* coding region we identified seven different mutations located in the transpeptidase domain. Three different mutation sites were identified independently in more than one lineage (Fig. S4B). Only one of the changes identified in this work coincided with previously described mutations (27), however we previously reported a mutation in position 617 of the protein in LS4828, which was correlated with decreased penicillin binding (9), and in this current work we identified a different change in the same position (Table S4). In addition to *pbp4* mutations we identified five lineages with mutations in the class A Pbp *ponA* gene, all mutations were within the putative transpeptidase domain (Fig. S4C), one of the three identified mutations occurred three times independently (Table S4).

The cell wall hydrolase N-acetylmuramoyl-L-alanine amidase (AmiA) has been implicated in septum digestion and peptidoglycan turnover (28). We identified mutations in the *amiA* gene in 6 independent lineages. Five of the six mutations were either premature stop codons or nucleotide deletions (Fig 4S, Table 4S), suggesting that the inactivation of this autolysin could help in peptidoglycan stability in the resistant mutants.

The enterococcal serine/threonine kinase and its cognate phosphatase (*stp*(*ireK*)/*stk*(*ireP*)) play a central role in resistance to antibiotics, mostly cephalosporins, and other cell wall stresses (29). Resistance is mediated by IreB phosphorylation. The lack of phosphorylation in IreB either by impairing substitutions or by the absence of IreK results in reduced cephalosporin resistance. To turn off the signaling pathway, the cognate phosphatase, IreP, dephosphorylates IreK (30, 31). We identified mutations in *ireP* and *ireB* in six of the sequenced resistant lineages (Table S4). Our data suggest that the identified mutations maintain IreB constantly phosphorylated or active, either by *ireP* inactivating mutations or by mutations that prevent IreB dephosphorylation.

The essential two component system (TCS) *walKR* controls cell wall metabolism in Firmicutes (32, 33) by the homeostatic regulation of the cell wall hydrolases required for peptidoglycan expansion during growth coordinating both processes (34). Mutations in the WalKR two component system were identified in 6 of 10 sequenced lineages. In the lineages derived from ATCC 29212, JH2-2, and D32, the mutations occurred in the sensor histidine kinase gene *walK*. The only sequenced lineage derived from a susceptible background that did not acquire *walK/R* mutations was JL1Amp (JH2-2) and this lineage did not become ampicillin resistant, suggesting that *walK* is important in the acquisition of ampicillin resistance. All WalK mutations were amino acid substitutions and the majority occurred early during the experiment. In LSL2Imi (LS4828), the mutation occurred in the *yycl* response regulator, which controls the activation/deactivation of the *walKR* system (Fig. S4D, Table 4S).

We identified mutations in other genes involved in cell wall homeostasis that were lineage or strain specific and not shared between the different resistant mutants as shown on the supplemental material (supplemental dataset).

### Mutations in the c-di-AMP pathway

We identified mutations in *cdaA, pgpH* and *gdpP* in our evolved lineages (Table S5). We identified *cdaA* mutations (amino acid substitutions) in three lineages (one derived from ATCC 29212 and two from JH2-2), the mutations were all located within the regions coding for the transmembrane domains. No mutations were identified within the catalytic deadenylate cyclase domain (Fig. S5A). The *cdaA* mutations occurred relatively early during the LTEE, between passages 14 and 70. We also identified a deletion in the putative *cdaA* regulator in one lineage derived from JH2-2. Additionally, we identified mutations in *pgpH* in five lineages from the four different backgrounds (Table S5) and *gdpP* mutations in two lineages derived from JH2-2. The mutations in the phosphodiesterases were either premature stop codons or amino acid substitutions in the catalytic domain (Fig. S5B, C, Table 5S).

### c-di-AMP is elevated in resistant and dependent lineages

Due to the ubiquity of mutations in c-di-AMP regulating genes during the selection process in the LTEE, we were interested in determining c-di-AMP levels at different time points in β-lactam resistant lineages. We performed c-di-AMP quantification by competitive ELISA. We found that *cdaA* mutations had little to no impact on c-di-AMP levels, however lineages with mutations in either *pgpH* or *gdpP* reached very high levels of intracellular c-di-AMP (Fig. 3). Trans complementation studies with a plasmid carrying a wild type copy of *pgpH* under a constitutive promoter significantly reduced c-di-AMP levels in two resistant and dependent lineages with *pgpH* mutations, improved growth and eliminated dependence without modifying the resistance to β-lactams (Fig. 4). Our results shown that all but one lineage with *pgpH* or *gdpP* mutations developed β-lactam dependence and that all dependent lineages sequenced carried *pgpH* or *gdpP* mutations, indicating that mutations in one of the two c-di-AMP degrading enzymes are necessary but not sufficient for the β-lactam dependent phenotype.

**Figure 3.**
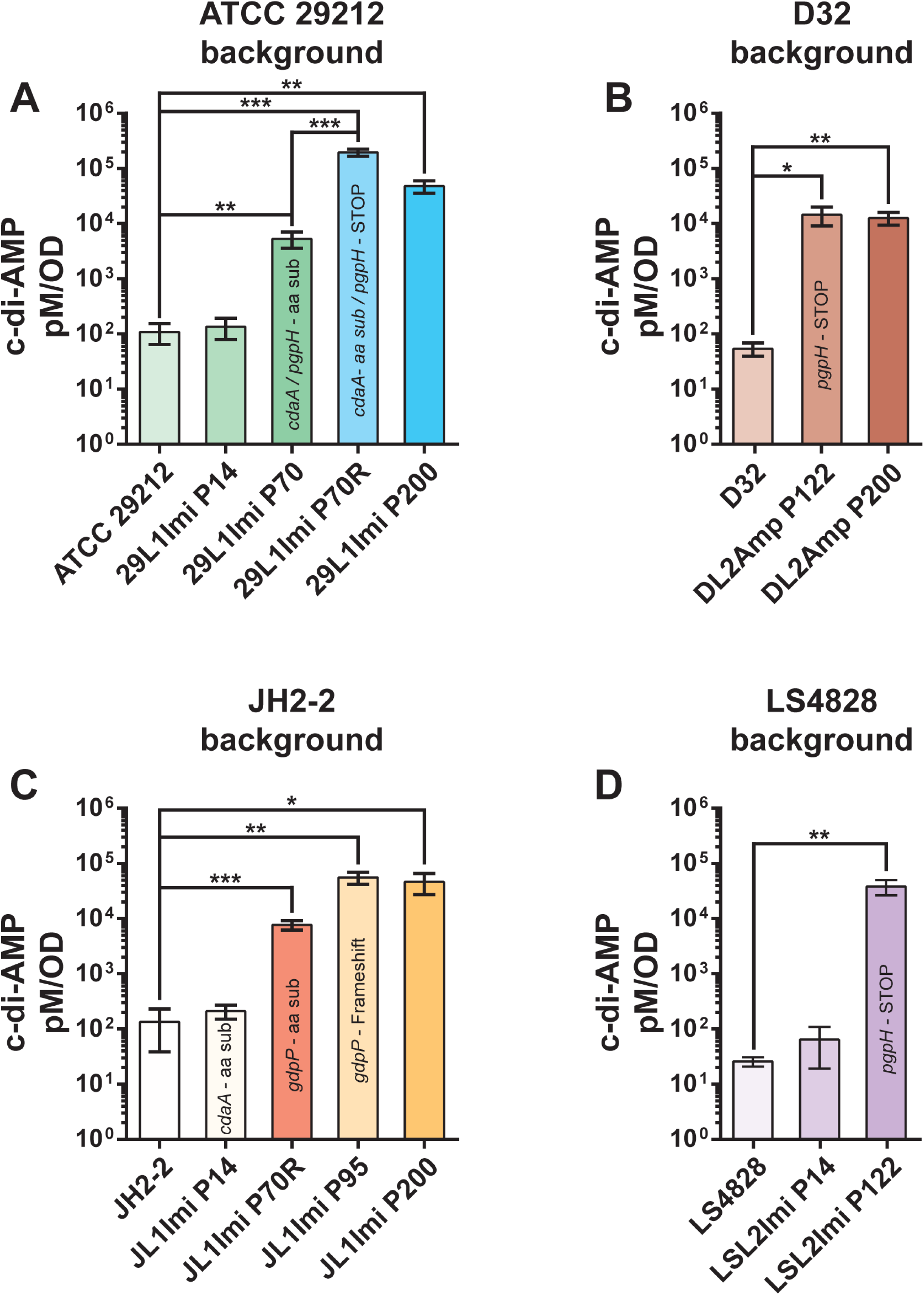
c-di-AMP quantification. The graphs show c-di-AMP levels in four different lineages carrying *cdaA* and or *pgpH/gdpP* mutations. The passage is which a particular mutation was detected is labeled with the mutated gene. **A)** ATCC 29212-derived lineages 29L1Imi **O**riginal and **R**eplica. *cdaA* and *pgpH* mutations were acquired before the split of the original and replica plates. **B)** D32-derived lineage DL2Amp, selected with ampicillin. This lineage acquired a inactivating pgpH mutation at P122 significantly increasing c-di-AMP concentrations. **C)** JH2-2-derived lineage JL1Imi, selected with imipenem. This lineage acquired a *cdaA* mutation at P14 that did not produce significant changes in c-di-AMP concentrations and independent *gdpP* mutations (P70R and P95O), that both correlate with a significant increase in c-di-AMP intracellular concentrations. D) LS4828-derived lineage LSL2Imi acquired an inactivating pgpH mutation at P122 with a significant increase in c-di-AMP levels. *p ≤ 0.05; **p ≤ 0.01.

**Figure 4.**
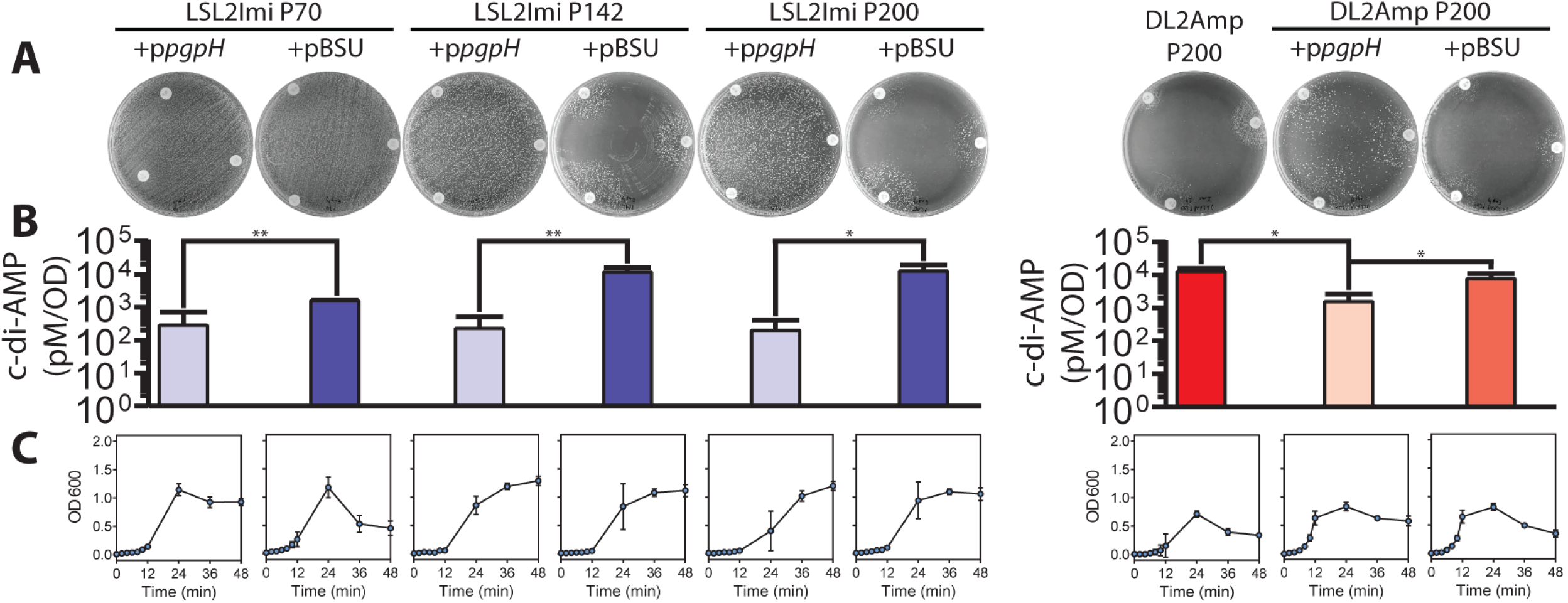
The presence of a functional copy of *pgpH* inhibits the β-lactam dependent phenotype by decreasing the level of c-di-AMP. Passages 70 (without *pgpH* mutation), 142, and 200 (with a premature STOP codon at *pgpH*) from lineage LSL2imi and passage 200 from DL2Amp, were transformed with the plasmid p*pgpH* (full-length *pgpH* with a constitutive promoter) or pBSU (negative control). **A)** Disc diffusion assay with the complemented passages using ampicillin (Bottom right of the plate,AM-10 µg), imipenem (Bottom left of the plate, IPM-10 µg), and penicillin (Top of the plate, P-10 µg) discs. The petri dishes show the complete inhibition (LSL2imi) or the softening (DL2Amp) of the β-lactam dependent phenotype when the passages express the full-length version of *pgpH*. **B)** C-di-AMP levels (pM/OD) in the complemented passages and their corresponding controls. Error bars indicate the standard deviation of the mean of three independent experiments. *: P ≤ 0.05; **: P ≤ 0.01. **C)** Growth profile of the complemented passages determined by optical density (OD600).

### Summary of genetic changes in β-lactam resistant lineages

The acquisition of β-lactam resistance documented throughout the adaptive evolution of our selected lineages suggests a role of distinct mutations that were acquired in more than one genome and that correlated with changes in the MICs. Other mutations may not exhibit a direct role in β-lactam resistance but may be adaptations to restore fitness. To gain a deeper insight into the evolutionary trajectory of β-lactam adaptation, we correlated various aspects observed throughout the evolution of four selected lineages. The aspects considered were increasing antibiotic concentration, MICs values, cumulative mutated genes, intracellular concentration of c-di-AMP, development of antibiotic dependence, and growth.

In the β-lactam resistant LS4828-derived lineage LSL2Imi, the first mutation identified was a premature stop codon in *mltg* at passage 37, eliminating the last 57 amino acids that corresponds to part of the putative active domain. *mltg* codes for a putative endolytic murein transglycosylase/peptidoglycan muramidase. We did not observe changes in the MIC’s at this time point, however we observed decreased fitness, determined as a delay in growth. Between passage 37 and 60, imipenem MIC increased from 8 µg/ml to >256 µg/ml and the lineage acquired mutations in *wbbl* and *rodA*. Neither *wbbl* nor *rodA* have been previously associated with antibiotic resistance in *Enterococcus*, but the role of RodA in cell wall homeostasis supports further studies. We observed cell lysis during stationary phase starting at passage 60 and until passage 122, lysis ended at passage 142 in association with a null mutation in the major autolysin gene *altA*. In passage 70 there was a single mutation in the WalK regulator gene *yycl*, we did not observe major changes in MICs or growth. Passage 95 had an increase of the ampicillin MIC to >256, however no new mutations were identified at this time point, suggesting that the cumulative effect of previous mutations help this lineage to withstand higher and higher ampicillin concentrations. The presence of a null mutation in the *pgpH* gene at passage 122 correlated with a statistically significant increase in intracellular c-di-AMP, compared to P95, and correlated with the emergence of β-lactam dependency (Fig. 5).

**Figure 5.**
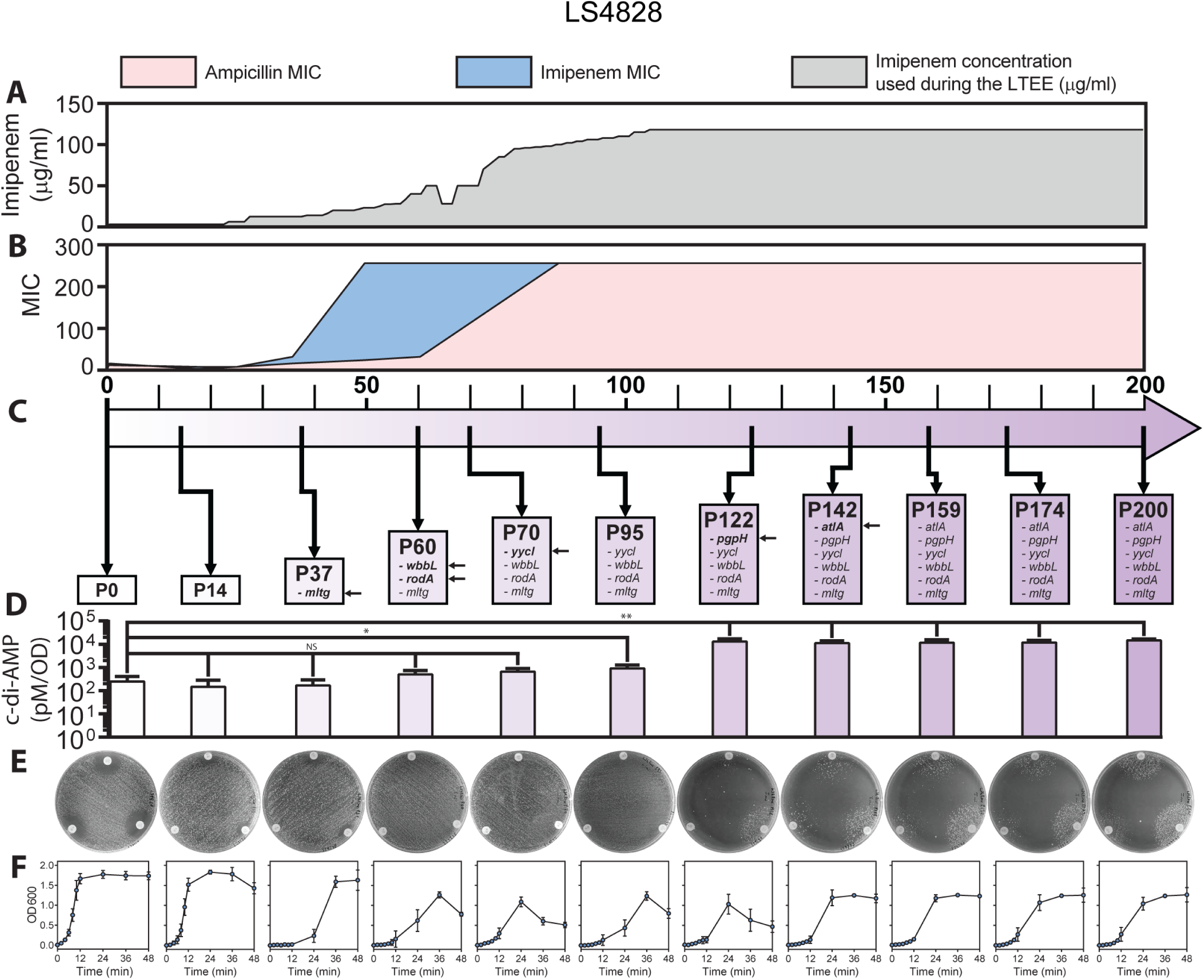
Timeline of the gradual adaptations to imipenem of LSL2Imi during the LTEE. **A)** gradual increment in imipenem concentration (µg/ml-grey) during the LTEE. **B)** Ampicillin (pink) and imipenem (blue) MICs. **C)** Mutated genes with a putative role in cell wall homeostasis (the squares correspond to the first time each mutation was identified. **D)** c-di-AMP levels (pM/OD), results from three independent experiments. A Student t-test applying a p>0.05 was used to compared passage 0 (parental strain) with all the rest of the passages. *: P ≤ 0.05; **: P ≤ 0.01; ***: P ≤ 0.001; ****, P ≤ 0.0001. **E)** Disc diffusion assay with ampicillin (Top of the plate, AM-10 µg), imipenem (Bottom right of the plate, IPM-10 µg), and penicillin (Bottom left of the plate, P-10 µg). The petri dishes show the transition from resistance to a β-lactam dependence along the experiment. **F)** Growth curves from three independent experiments.

In JH2-2-derived lineage JL1Imi, the first two mutations were detected early during the experiment, when the imipenem concentration used for selection was still very low, at passage 14 in *cdaA* and *bgsB* (1,2-diacylglycerol 3-glucosyltransferase), correlating with a delay in growth. Growth did not recover in this lineage, and the growth curves show growth only after 24h or later, suggesting that the population recovery observed is due to new mutations acquired during the growth curve experiment (Fig. 6F), we currently do not have sequencing data for this samples. Mutations in *walK* and *lpa* (lipoamidase) at passages 60 and 70, respectively, were associated with a moderate increase in MIC for ampicillin and imipenem. At passage 70 we observed a sharp decrease in fitness, measured as growth. As we progressively increased the imipenem concentration, we documented *ireB* and *gdpP* mutations at passage 95 in the original culture (Fig.6A-F), these mutations coincided with the development of β-lactam dependence and an inability to obtain MIC readings at 24h due to delayed growth. The presence of a mutation in *lepB* (a signal peptidase) at passage 142 correlated with better growth. This lineage acquired a *pbp4* mutation later in the experiment at passage 174.

**Figure 6.**
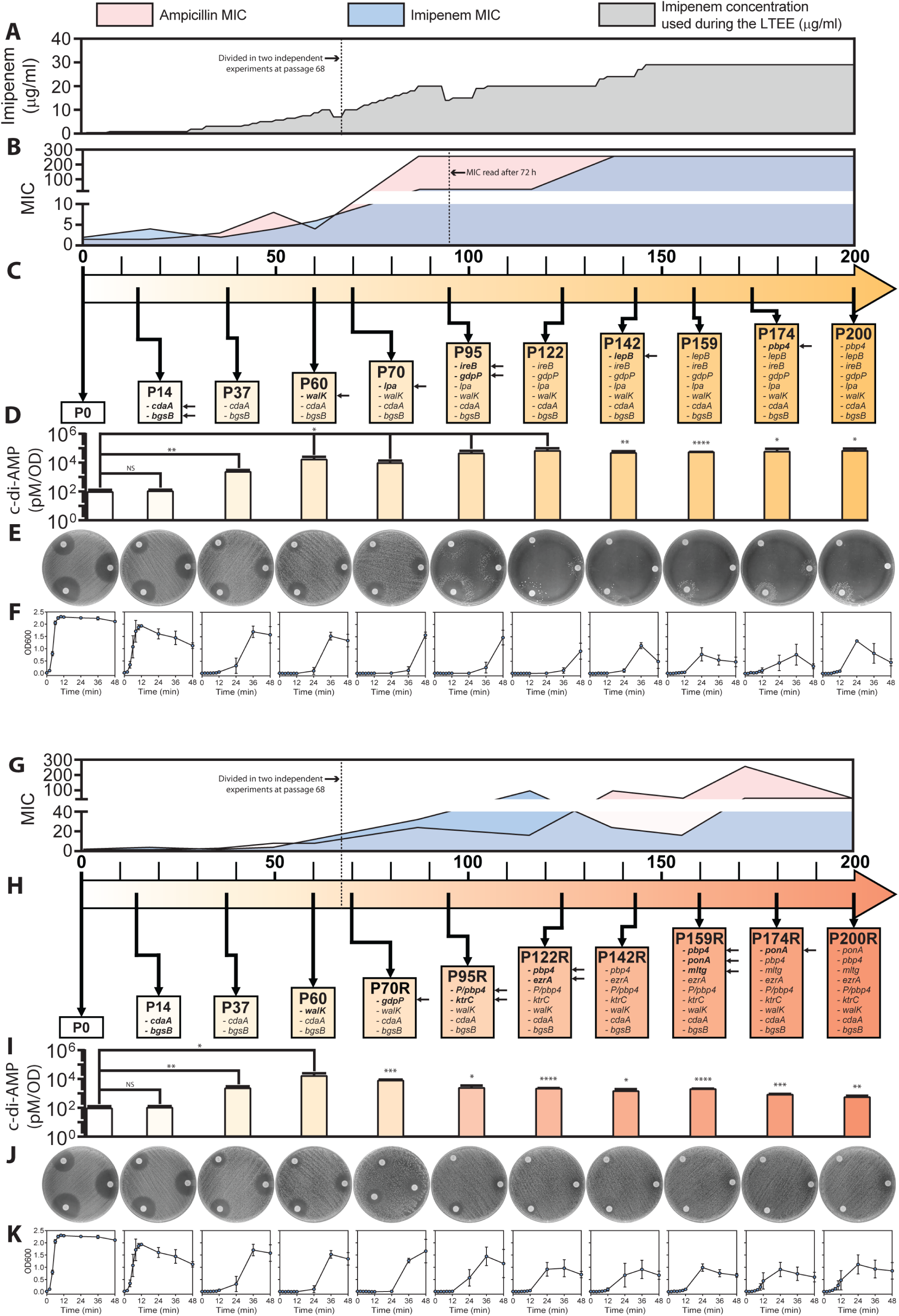
Timeline of the gradual adaptations to imipenem of JL1Imi during the LTEE in original and replica cultures. **A)** gradual increment in the imipenem concentration (µg/ml-grey) during the LTEE. **B)** MICs of ampicillin (pink) and imipenem (blue). The MICs were read from passage 95 and up to 200 at 72 h because of the growing delay of those passages in the original culture. **C)** Mutated genes with a putative role in resistance or fitness restoration. The mutated genes that appeared for the first time in specific passages are pointed out by arrows. **D)** Levels of c-di-AMP (pM/OD) throughout the experiment. Error bars indicate the standard deviation of the mean of three independent experiments. A Student t-test applying a p>0.05 was used to compared passage 0 (parental strain) with all the rest of the passages. *: P ≤ 0.05; **: P ≤ 0.01; ***: P ≤ 0.001; ****, P ≤ 0.0001. **E)** Disc diffusion assay with ampicillin (Bottom right of the plate, AM-10 µg), imipenem (Bottom left of the plate, IPM-10 µg), and penicillin (Top left of the plate, P-10 µg). The petri dishes show the transition of the passages from a resistant phenotype to a β-lactam dependent phenotype across the experiment. **F)** Growth curves determined by optical density (OD600). The lines in the graphs represent the mean of three independent experiments. Error bars indicate the standard deviation. The same aspects were measured in the replica culture. **G)** MICs. **H)** Mutated genes. **I)** c-di-AMP levels. **J)** Disc diffusion assay. **K)** Growth curves.

The replica culture (Fig. 6G-K) once split from the original culture, evolved a different trajectory. At passage 70 it acquired an independent mutation in *gdpP* (amino acid substitution, compared to premature stop codon), which coincided with a modest increase in MIC and significant increase in c-di-AMP concentrations compared to previous passages, however c-di-AMP levels were not as high as in lineages carrying premature stop codons. Interestingly this lineage did not develop dependence. Additionally, this lineage acquired four independent mutations in *pbp4* (detected at passages 95R, 122R, 159R and 200R). Ampicillin MICs increased at passages 95R, 142-149R and 174R. Imipenem MICs increased at passages 95R and 122R. In addition to *pbp4* mutations we identified two mutations in *ponA* (passages 159R and 174R), and a convergent mutation in *lpa*.

In the ATCC 29212-derived lineages 29L1ImiO and 29L1ImiR followed different trajectories to acquire β-lactam resistance and to develop antibiotic dependence. 29L1ImiR lineage acquired resistance to ampicillin and imipenem at passage 60 and passage 70R respectively, at this time we identified mutations in *cdaA, pgpH, pbp4, ireB* and *walK* and we observed an initially weak dependent phenotype at passage 70R. A mutation in *ponA* at passage 95R coincided with a further increase in MICs. Two additional mutations in *pbp4* (passages 122R and 174R) and a mutation in *amiA* (passage 174R) seemed to not have a further impact on the MICs, however MIC testing in this lineage show fluctuating results and high variability.

The dependent phenotype is highly variable and does not always correlate temporarily with the acquisition of mutations in the c-di-AMP pathway or with the peak concentration of this metabolite, as exemplified in the differences observed between the original and replica cultures of 29L1Imi. 29L1ImiO acquired mutations in *cdaA* and *pgpH* (independently of 29L1ImiR) at passage 70 but developed a dependent phenotype only at passage 95 and later lost dependency at passage 175. DL2Amp was derived from D32 under selection with ampicillin. Resistance to ampicillin was acquired at passage 95 when it had accumulated *walK, pbp2B,* and *ireP* mutations, resistance to imipenem was detected at passage 122, coincidental with a *pgpH* mutation and increased c-di-AMP levels. A β-lactam dependent phenotype was evident only in passages 174 and 200.

Mutations in *pgpH* and elevated levels of c-di-AMP seem to be necessary for the development of β-lactam dependence.

## Discussion

The acquisition of β-lactam resistance is a complex phenomenon, accompanied by evolutionary trade-offs. The results presented in this paper confirm the capacity of *E. faecalis* strains to acquire markedly elevated levels of resistance when confronted with persistent selective pressure from β-lactam antibiotics. We were able to achieve levels of resistance considerably greater than obtained in previous reports (9, 35). The use of four distinct genetic backgrounds and the splitting of these strains into different lineages allowed us to look for convergent genes and pathways involved in resistance, of which we found several. While clinical isolates demonstrating MICs to ampicillin and imipenem ranging between 12 and 32 µg/ml have been documented (9, 35), high MICs of 256 µg/ml to both ampicillin and imipenem are uncommon among *E. faecalis* isolates. Nevertheless, there is a report of some *E. faecalis* isolates from patients that reached MIC for imipenem of >256 µg/ml, although the genetic or metabolic determinants involved remain elusive (36). We did not identify additional reports of either clinical isolates or laboratory evolved lineages of *E. faecalis* achieving the higher MICs observed in our study (>256 µg/ml). Enterococci typically display resistance to most β-lactams, exhibiting relative susceptibilities to certain members of the penicillin group such as ampicillin, penicillin, and piperacillin(1). Resistance to ampicillin and imipenem in *E. faecalis* is closely linked to the overexpression of low-affinity class B Pbp4 induced by mutations in the promoter region. We and others had previously reported promoter deletions leading to *pbp4* overexpression in both clinical and laboratory strains (9, 37–39), suggesting that the mutations identified in the current work will have a similar effect in *pbp4* transcription. Amino acid substitutions have also been implicated in ampicillin/imipenem resistance in *E. faecalis* (9, 35, 38, 40–42), but the resulting resistance levels are not as pronounced as those reported here. Hence, we consider that the exceptionally high resistance levels observed are attributed to the sum of factors in addition to Pbp4.

Our data indicate that resistance in virtually all strains emerges more easily to imipenem than to ampicillin, and it emerges more readily under imipenem selective pressure than under pressure from ampicillin, especially in the strains that were initially susceptible. These results are in line of what we and others have reported with the clinical use of amoxicillin or imipenem (9, 37). To our knowledge there are not known differences in the effects of imipenem versus ampicillin on *E. faecalis* cells, such as differential reactivities of imipenem and ampicillin with the PBPs that could explain the differences observed in our study. Further work would provide interesting insights into this observation.

As mentioned before, β-lactam resistance in *E. faecalis* has most frequently been tied to changes within Pbp4. These changes have additionally involved amino acid substitutions, many of which are assumed to reduce affinity for the antibiotic, but only a few of which have been documented to do so (27, 35, 38, 43). We observed seven different amino acid substitutions in Pbp4 over the course of the experiment, with two appearing in more than one strain. While we do not have experimental evidence that these changes confer resistance, they are all in proximity to the active site (personal communication, Wolfgang Peti), so may either alone or in combination confer reduced susceptibility to ampicillin, imipenem, or both.

It is conceivable that prolonged and consistent exposure to β-lactams may induce a recurring pattern of mutations within specific genes associated with resistance. It is also noteworthy that strain LS4828, a strain clinically resistant to β-lactams underwent no further Ppb4 substitutions yet achieved high levels of resistance in the shortest time compared with the other three susceptible backgrounds. The increase of ampicillin and imipenem MICs in LS4828-evolved lineages cannot be attributed to *pbp4* mutations. One gene that is interesting in the context of β-lactam resistance is *mltG*, coding for a putative peptidoglycan muramidase essential for cell elongation in streptococci (44). Additional mutation occurs in RodA, a glycosytransferase that partners with PBP transpeptidases to make peptidoglycan during cellular elongation and division (45), and WbbL (L-rhamnosyltransferase of unknown function in *E. faecalis* that has analogues in Gram-negative bacteria that are involved in lipopolysaccharide biosynthesis and is important for synthesis of O-antigen) (46), could be contributing to further increases in the imipenem MICs. Although the interaction between MltG and RodA have not been investigated in *E. faecalis*, this interaction was reported in *E. coli* (47). Additionally, a model of how these proteins interact was proposed in *S. pneumoniae*, in which MltG and other amidases open the peptidoglycan layer, letting RodA to interact with a PBP to add new peptidoglycans to the layer, inducing the elongation of the cell (45). Other interesting gene is *yycl*, a regulator of the WalR/WalK two component system, which has been shown to be involved in cell wall synthesis and antimicrobial susceptibility in different species, including *Staphylococcus aureus* and *Bacillus subtilis* (34, 48–50). In *S. aureus* the disruption of *yycH* and *yycI* genes led to a downregulation of the WalKR regulon, including reduced expression of the autolysin genes *atlA* and *sle1* inducing impaired cell wall turnover and reduced vancomycin susceptibility (51). In contrast, in *B. subtilis* cells lacking *yycH* or *yycl* showed stronger transcription of autolysins, suggesting that the active genes lead to a reduction of WalK activity (52–54). Our data suggest that in *E. faecalis* the disruption of *yycl* induces an activation of the WalKR regulon and that compensatory mutations such as the frameshift observed in *atlA* gene compensate an overactive WalKR and help keep cell wall turnover in check, improving growth rate at 12h. Further studies are necessary to validate the role of the mutated genes.

Our data indicate that there is strain specificity in the types of phenotypic resistance and genetic changes observed when the strains are under selective antimicrobial pressure. The pattern of mutations observed on different strains selected under ampicillin or imipenem pressure are more similar within the strains than between strains. A similar observation was made by Card and colleagues in *E. coli* (21), who concluded that strain genotype can affect both the genotypic and phenotypic pathways to resistance.

An intriguing finding of our study is that there appears to be a final common phenotypic characteristic of all strains – antibiotic dependence. This is a lethal phenotype in the absence of antibiotics, however eventually progeny emerge that can re-gain growth in the absence of antibiotics. Loss of dependence does not affect β-lactam resistance. The evolutionary advantage of β-lactam dependence vs. β-lactam resistance could rely in the inactivation by the antibiotic of other PBPs letting a highly resistant PBP4 take over the synthesis of cell wall. β-lactam dependent-lineages do not overexpress PBP4 upon exposure to the antibiotics. The evolution of β-lactam dependence in our selected lineages was associated with inactivating mutations in one of the two genes coding for the c-di-AMP phosphodiesterases. Consistent with the inactivating nature of these mutations, we were able to measure markedly increased quantities of c-di-AMP in mutant lineages. We confirmed the involvement of c-di-AMP levels in this phenotype by introducing a plasmid-encoded intact *pgpH*, maintaining β-lactam resistance and losing the dependent phenotype. In *Staphylococcus aureus* inactivating mutations have been associated with β-lactam tolerance and enhanced evolution of resistance both in vitro and in clinical strains (55, 56). Tolerance to β-lactams could be caused by a modulation in cell wall thickness (57), further studies are necessary to determine if a similar mechanism occurs in *E. faecalis* and how this leads to β-lactam dependence. C-di-AMP is an important second messenger in gram-positive bacteria (58) and it has been implicated in virulence, susceptibility to cell envelope-targeting drugs, cell membrane acting antibiotics, peptide and metabolite transport, osmotic regulation and other fundamental cell functions (59). Homeostasis of c-di-AMP is of paramount importance since under many growth conditions it is essential, but its accumulation is also toxic (60–62). *E. faecalis* codes for a single diadenylate cyclase (CdaA), which is responsible for c-di-AMP synthesis, and two phosphodiesterases in charge of c-di-AMP degradation (PgpH and GdpP) (58, 60–64). C-di-AMP is essential during normal growth conditions in *E. faecalis* (64).

While some mutations are clearly associated with resistance or dependence, it is plausible that others are more closely linked with fitness restoration after the tradeoff of antibiotic resistance. Mutations improving fitness of resistant mutants can be epistatic, jointly contributing to adaptation under specific pressures (65). However, owing to the inherently stochastic dynamics of β-lactam resistance and fitness-related epistatic interactions, the data gathered in this study represents just a few of many possible evolutionary trajectories. This data may contribute towards uncovering patterns for predicting models that closely mirror the trajectory of *E. faecalis* evolution in response to β-lactam antibiotics. Consequently, more focused studies will be necessary in the future.

## Materials and Methods

### Long-term evolution experiment

To obtain β-lactam resistant lineages, four *E. faecalis* strains from different origins and with different initial susceptibilities (Table 1) were subjected to serial passaging in a LTEE. We exposed the four distinct *E. faecalis* strains to continuous and increasing ampicillin or imipenem selective pressure for a total of 6,000 generations (200 days). We started the LTEE from 3 founding colonies from each strain. Each founding colony was seeded into ampicillin, imipenem or no antibiotic, establishing a total of 36 initial cultures: 12 were grown with ampicillin, 12 were grown with imipenem, and 12 were grown in broth alone. The antibiotic concentrations were increased gradually depending on growth rate until the experiment reached 104 daily passages. After day 105, we kept the antibiotic concentration constant until day 200 because several lineages displayed very low growth rates. Samples were frozen every ten passages for each culture (Fig. S1). The initial ampicillin and imipenem concentration was ¼ MIC for each strain. The LTEE was carried out in BHI media. Cultures were passaged by 100-fold dilution (5 µl in 500 µl) every 24h as reported by Lenski and coworkers in 1994 (76). The initial inoculum at the start of the experiment was OD_600_ ∼0.001 (∼7.00E+03 CFU’s). Initially, when the antibiotic selection presure was low, growth after 24h in all plates was about 7.70E+09 CFU’s/ml (OD_600_ 0.8), and each fresh well was inoculated with 7.70E+07 CFUs/ml (∼3.85E+05 in 5µl). As the LTEE progressed and the growth rate became more variable between the different lineages the CFU’s inoculated were also more variable, moreover, considering that several lineages developed a clumpy phenotype that made accurate quantification complicated. Weekly sampling and storage were performed for each lineage. MICs to ampicillin and imipenem were determined every ∼450 generations (15 days) by MIC strip tests on MHII agar plates.

### Genomic analysis

Whole genome sequencing was performed by Psomagen (MD) or SeqCenter (PA). Read length was 151 reads with average coverage of 2000X (Psomagen) and 30X (SeqCenter). De novo genome assembly was performed with SPAdes version 3.12 (79), using the default parameters. Genome annotation was performed with Prokka (80). SPAdes and Prokka were run through the Brown University supercomputing cluster OSCAR. Variant calling was performed with Geneious 11.1.5 using default parameters for bacterial genomes, additional variant calling analysis was carried out by SeqCenter using breseq version 0.35.4 (81).

### Intracellular cyclic di-AMP measurement by Competitive ELISA

Cyclic di-AMP ELISA (Enzyme-Linked Immunosorbent Assay) Kit (Cayman Chemical) was used for quantification of c-di-AMP. Briefly, the samples were assayed using a minimum of two dilutions in duplicate. Sample and standard volumes of 50 µl were added to the pre-coated goat anti-mouse IgG and pre-blocked microtiter plate followed by the addition of 50 µl of Cyclic di-AMP-HRP (horseradish peroxidase conjugate) Tracer and 50 µl of Cyclic di-AMP Monoclonal Antibody. Appropriate controls according to the manufacturer’s directions were also added at this point. The plate was covered and incubated at room temperature on an orbital shaker for 2 h. The wells were emptied, washed 5 times with provided Wash Buffer with Tween 20, and filled with 175 µl of TMB (3,3’,5,5’-Tetramethylbenzidine) Substrate Solution plus 5 µl of the diluted tracer to the appropriate well. The plate was covered and incubated at room temperature for 30 min on the orbital shaker. The reaction was stopped with 75 µl of HRP Stop Solution and the plate was read at 450 nm in a BioTek EPOCH2 microplate reader. The data were analyzed using a computer spreadsheet provided by Cayman Chemical. Using the spreadsheet, the standard curve data was linearized using a logit transformation and the sample values were determined from the standard curve.

### Statistical analysis

Data from c-di-AMP measurements were analyzed with GraphPad Prism version 6.0 (GraphPad Software, San Diego, CA, USA). Data from three independent experiments were plotted and a two-tailed Student t-test was used to compare each group versus each other. A p>0.05 value was considered statistically significant.

## Data Availability

WGS data is available from NCBI with accession numbers: CP166607, JBGCWG000000000, JBGCWH000000000, JBGCWF000000000. Variant Calling Analysis is available as supplemental datasets 1,2,3 and 4. Protein accession numbers are provided in Table S6

## Author contributions

PUS, investigation, methodology, formal analysis, visualization, writing-original draft; CD: investigation, methodology, writing-review and editing; LBR: resources, funding acquisition, writing-review and editing; MGS: conceptualization, conceived and planned the project, data curation, formal analysis, investigation, methodology, project administration, supervision, validation, writing-review and editing.

## Acknowledgments

**This work was supported in part by R01AI141522-05 from the National Institute of Allergy and Infectious Diseases (L.B.R.).** The authors thank Wofgang Peti for modelling Pbp4 mutations; Juan Camillo Carrillo Martinez for his help performing an additional growth curve for figure 6.

## Supplemental Material

### Supplemental Methods

#### Bacterial Strains

We used four different *E. faecalis* strains for which whole genome information was available and were isolated from diverse sources (Table 1): D32 (25) lacks distinct virulence-associated traits (66). D32 belongs to ST40; strains from this sequence type do not show ecological preference and have been isolated from animals, food, and humans (67). ATCC 29212 (68, 69), it belongs to ST30, which is associated with humans and dates back to at least the 1950s (67, 70) and contains several described virulence factors (69). JH2-2 (24), has been long used as a laboratory strain. LS4828 is an ampicillin resistant strain that was isolated from a prosthetic knee after prolonged treatment with amoxicillin. LS4828 overexpresses *pbp4* and the PBP4 protein has reduced affinity to β-lactams (9).

#### Antimicrobials

Ampicillin (Sigma Aldrich, St. Louis, MO), Imipenem (Combi-Blocks, San Diego, CA), MIC test strips were acquired from Liofilchem: ampicillin 0.016-256 µg/ml, ampicillin 0.002-32 µg/ml, imipenem 0.016-256 µg/ml, imipenem 0.002-32 µg/ml, penicillin 0.016-256 µg/ml, penicillin 0.002-32 µg/ml. Antibiotic discs were acquired from BD BBLTM.

#### Media

As per Clinical and Laboratory Standards Institute (CLSI) guidance, Mueller-Hinton II broth (MHII; Becton Dickinson, Sparks, MD, USA), adjusted to 25 mg/L calcium and 12.5 mg/L magnesium, was used for all susceptibility testing as previously described (71–74). Liquid culture for the LTEE and streaks were grown with brain heart infusion (BHI) (Sigma Aldrich, St. Louis, MO) due to better growth (9, 75, 76). For colony counts, MHII agar plates were used for better resolution to count.

#### Minimal inhibitory concentration (MIC) testing

10 µl from each sample was diluted in 3 ml of BHI broth. The entire surface of regular BHI agar Petri plate was inoculated in three different directions by using cotton swabs dipped in the inoculum suspensions. MIC strip tests were applied onto the inoculated agar using forceps. The plates were incubated at 37° C and read by eye after 18-24 h of incubation.

#### Disc diffusion assays

Overnight cultures were adjusted to OD_600_ 0.01 of in 3 ml of BHI broth supplemented with either ampicillin or imipenem, depending on the antibiotic selection and concentrations used during the LTEE. The cultures were incubated until they reached OD_600_ an ∼0.5. The OD_600_ was adjusted to 0.01. The entire surface of regular BHI agar Petri plate was inoculated in three different directions by using cotton swabs dipped in the inoculum suspensions. Disks were applied to the inoculated agar by using forceps. The plates were incubated at 37° C and the inhibition zone was measured after 24, 48,72 and 96 h.

#### Growth Curves

Lineages were streaked fresh from -80° C stored glycerol stocks onto BHI agar Petri plates. An ampicillin or imipenem disc was placed on the center of the agar plate. Additionally, complemented lineages were inoculated onto BHI agar Petri plates supplemented with spectinomycin at 300 µg/ml. The plates were incubated for 18 h at 37° C (or longer in the case of slow-growing dependent lineages). Three to five colonies were picked up near the disc and inoculated into BHI broth supplemented with either ampicillin or imipenem. Cultures were incubated at 37° C overnight or longer depending on growth rate. Fresh 20 ml of BHI supplemented with the corresponding β-lactam were inoculated at an OD_600_ 0.01 and incubated in a shaking incubator at 37° C and 180 rpm. Growth was monitored by OD_600_ at 0, 4, 8, 12, 24, 36, and 48 h.

For CFU counts, overnight cultures were washed three times with saline solution to clean remains of antibiotic used for growth overnight. Cultures were inoculated at an initial OD_600_ 0.01 into 5 ml of fresh BHI broth supplemented with or without the corresponding β-lactam. The cultures were incubated in a shaking incubator at 37° C and 180 rpm. Samples were obtained at 0, 4, 8, 12, 24, 36, and 48 h. Samples were serially diluted and plated onto MHI agar plates to count and calculate the CFU per ml. The plates were supplemented with 5 µg/ml of ampicillin for β-lactam dependent lineages. The lower limit of detection (LLD) was 2.0 log10 CFU/ml.

### Plasmid construction and transformation

The coding region of LS4828 *pgpH* was cloned by PCR and restriction digestion into the BamHI and XbaI sites of pBSU101 (77), producing a fusion of the pgpH CDS with the plasmid CFB promoter. The sequences of the plasmid were confirmed, and they were transformed into *E. faecalis* by electroporation.

Electrocompetent cells were prepared according to the protocol of Friesenegger et al. 1991 (78) with modifications. Cells were grown in Todd-Hewitt broth supplemented with the corresponding antibiotic used during the LTEE. Overnight cultures were diluted 1:1000 in 150 mL of Todd-Hewitt broth and incubated for 12 hours or until stationary phase was reached. Cells were harvested by centrifugation at 3800 x g for 15 minutes and washed in cold 10% glycerol with 1/1, ½, ¼ and 1/8 of the initial culture volume. The cell pellets were resuspended in 1 mL of 10% glycerol, aliquoted, and frozen at -80°C. Electroporation was achieved by following the protocol of (78). Fifty microliter aliquots of electrocompetent cells were thawed on ice and mixed with ∼500 ng of pBSU101 or *p*pgpH DNA. Mixes were transferred to an ice-cooled 0.2 cm cuvette and pulsed with field strength of 12.500 V/cm using the 400Ω resistor to result in time constants of 9-16 ms. Cells were recovered in 1 mL Todd-Hewitt broth for 1 hr. at 37°C with 180 rpm shaking. One-hundred or 300 microliters of cells were spread onto BHI agar plates supplemented with 300 µg/mL spectinomycin and 5 µg/mL ampicillin. The plates were incubated for 48 hr. at 37°C to allow growth of transformants for selection. Colonies were passed one time to a fresh BHI agar plate plus spectinomycin 150 ug/mL and an ampicillin disc (10 µg) placed centrally prior to growing the transformants in broth for glycerol stocks and experiments.

### Genomic extraction

Genomic DNA was extracted from 2 mL of culture at mid log phase (OD_600_ of 0.500) using the Monarch Genomic Minikit (New England Biolabs, MA), following the manufacturer’s protocol with the following modifications: cells were lysed with 10 mg/mL lysozyme in Tris-EDTA pH 8 for 1 hour at 37°C. The DNA library was prepared using the TruSeq DNA PCR free kit 350 (Illumina) following the manufacturer’s user guide. The initial concentration of DNA was evaluated using the Qubit® dsDNA HS Assay Kit (Life Technologies), and 50 ng DNA was used to prepare the library.

### Preparation of samples for c-di-AMP Competitive ELISA

Parental strains and evolved lineages were grown and prepared for intracellular cyclic di-AMP measurements with some modifications according to the protocol of Wang et al. 2017 (61). Parental strains and evolved lineages were inoculated from glycerol stocks into 3 ml of regular BHI broth or BHI supplemented with ampicillin or imipenem, depending on the antibiotic or concentration used during the LTEE, and incubated with shaking at 180 rpm at 37° C for 17-48 h, depending on the strain, passage, or lineage. After incubation, cultures were diluted back 1:50 or 1:100 into 55 mL BHI supplemented with the corresponding antibiotic and grown with shaking at 180 rpm at 37° C to late exponential phase (OD_600_ ∼1.0). A minimum of 3 independent biological replicates were obtained for analysis.

Cell pellets from 45 ml culture volumes were centrifuged at 4000 g for 20 min and washed 3 times with 5 ml phosphate-buffered saline (PBS) followed by centrifugation at 4000 g for 5 min., pouring off PBS and freezing the cell pellet at -20oC. The cells were re-suspended in 0.5 ml of 50 mM Tris-HCL pH 8.0 and broken using Lysing Matrix B tubes (Thermo Fisher Scientific, Waltham, MA) with a BioSpec Mini Beadbeater-16 bead for 20 sec five times, with 1 minute on ice in-between. Cell lysates were collected by centrifuging for 15 min at 12,000 g. The lysate was transferred to a new tube, removing a small sample for protein concentration (Pierce BCA protein assay kit; Thermo Fisher Scientific). Protein concentrations were used to confirm complete cell lysis of samples. The remaining supernatant was boiled at 95° C for 5 min, cooled, and collected by centrifugation at 12,000 g for 15 min at 4° C. The protein-free supernatants were stored at -80° C until run in the competitive ELISA specific for c-di-AMP.

## Supplemental Tables

**Supplemental Table S1. Lineage identification key and sequenced genomes.**

**Supplemental Table S2. Ampicillin and imipenem MICs from lineages with no selective pressure.** Heat map of the Minimal Inhibitory Concentration (MIC) of ampicillin (red) and imipenem (blue) obtained from the lineages evolved without antibiotic selective pressure. MICs were determined between 18-24h of incubation and read by eye. The vertical dotted line delimits the passages from the original culture and the passages from the replica culture.

**Supplemental Table S3. Disc diffusion assays.** Disc diffusion assay in dependent lineages at passage 200 to determine resistance/susceptibility and dependence. β-lactam dependency is observed with different groups of β-lactams. Disc diffusion assays with penicillin (P-10 µg), oxacillin (OX-1 µg), ticarcillin (TIC-75 µg), piperacillin (PRL-100 µg), ampicillin (AM-10 µg), imipenem (IPM-10 µg), cefazolin (KZ-30 µg), cefuroxime (CXM-30 µg), cefotetan (CTT-30 µg), ceftazidime (CAZ-30 µg), ceftriaxone (CRO-30 µg), cefaclor (CEC-30 µg), cefepime (FEP-30 µg), and vancomycin (VA-30 µg). Mean diameter for three independent experiments. Cells were plated onto BHI agar Petri dishes at an OD600 of 0.01. The diameter of the growth or inhibitory halos were measured at 24, 48, 72, and 96 h.

**Supplemental Table S4. Mutations identified in genes related to cell wall synthesis and degradation.**

**Supplemental Table S5. Mutations in genes of the c-di-AMP biosynthetic pathway.**

**Supplemental Table S6. Protein accession numbers.**

## Supplemental Figures

**Supplemental Figure 1. Set up of the long-term evolutionary experiment (LTEE). A)** Three single colonies from each strain were picked up from BHI agar plates and seeded into three independent cultures (5 ml of BHI broth) establishing three lineages per strain (L1, L2, and L3) as biological replicas. **B)** Each founding lineage was split into three conditions: ampicillin (Amp), imipenem (Imi), or no treatment (NT). **C)** Plates were incubated at 37°C with shaking at 180 rpm for 24 h. Serial passages were performed daily by transferring 5 µl of each well into 500 µl of fresh BHI with the corresponding antibiotic concentrations. The experiment consisted of 200 consecutive daily passages. Amp or Imi concentrations were increased gradually up to day 104, and then it remained constant until day 200. **D)** Periodic freezing of samples and determination of MICs were performed. Amp and Imi MICs were determined by MIC strip test. **E)** Genomic DNA was obtained from specific samples to perform whole genome sequencing and variant calling.

**Supplemental figure 2. β-lactam dependent lineages grow dynamics.** The dependent lines grew faster in the presence of β-lactams. **A)** Growth curves with CFU/ml count of four β-lactam dependent lineages from passage 200 (P200): LSL2Imi P200OR (LS4828 background, original culture), 29L1Imi P200NRP (ATCC29212, replica culture), JL1Imi P200OR (JH2-2 background, original culture), and DL2Amp P200 (D32 background, original culture). The cells were grown overnight (ON) in the presence of ampicillin or imipenem. The ON cultures were washed three times with saline solution and adjusted to an initial 0.01 OD_600_ in BHI broth. The adjusted cultures were grown in the presence or absence of either ampicillin or imipenem using 118 ug/ml for LSL2imi, and 29 ug/ml for the other backgrounds, the same concentrations were used during the LTEE for those specific lineages. Samples were taken at 0, 4, 8, 12, 24, 48, and 72 h for CFU counts. The upper left area of the graph highlights the exponential growth phase in the presence of β-lactams. The lower left shows the population decline in the absence of antibiotic. The right side of the graph highlights the lag in exponential growth phase of the cultures without antibiotic and the stationary phase. **B)** Disc diffusion assay with ampicillin (Right side of the plate, AM-10 µg), imipenem (Bottom left of the plate, IPM-10 µg), and penicillin (Top left of the plate, P-10 µg). Cells were grown in BHI broth with or without the antibiotic in which they were originally selected until an 0.5 OD_600_. The cells were diluted to an 0.01 OD_600_ and streaked onto BHI agar Petri dishes. LSL2Imi P200 and DL2Amp P200 grew throughout the plate without a clearing inhibition zone. Lineage 29L1Imi P200R had inhibitory halos of 1.3 ± 0.57 mm to ampicillin, 4 ± 1 mmm to imipenem, and penicillin resistance, lineage JL1Imi P200 had inhibitory halos of 10.3 ± 0.57 mm to ampicillin, 9.6 ± 1.5 mmm to imipenem, and 7.3 ± 1.1 mm to penicillin. An inhibition zone diameter ≥ 15 mm indicates penicillin susceptibility and ≥ 17 mm indicates susceptibility to other β-lactams.

**Supplemental figure S3. Quantity and type of mutations detected in sequenced evolved lineages.** Summary of the number and type of mutations accumulated over time by seven selected lineages. **A)** Total number of mutations acquired by the lineages in the original (dark blue) and replica (light blue) culture. The mutations were grouped into five categories depending on the genetic change: deletions >50 bp, indel ≤50 bp, intergenic, non-sense, and non-synonymous. The number of mutations per category was counted in the genomes of **A)** LSL3Amp, **C)** 29L1Amp, **D)** JL1Amp, **E)** DL2Amp, **F)** LSL2Imi, **G)** 29L1Imi, and **H)** JL1Imi.

**Supplemental figure S4. Mutations in genes related to β-lactam resistance and cell wall regulation identified in more than one resistant lineage. A)** *pbp4* promoter region with the detected mutations, positions are relative to the gene start codon. **B)** Pbp4 domain organization and mutated residues. **C)** PonA mutations. **D)** WalK mutations. **E)** Mltg mutations. **F)** IreB mutations, LS4828-*ireB* has an insertion of an A at position 51 of the CDS, causing a frameshift in the protein. **G)** IreP mutations. Squares express the alignment of the mutated regions from the four different genetic backgrounds. The corresponding mutations are highlighted in red. Asterisks srepresent premature STOP codons. The mutated base or amino acids are bolded and squared within the alignment. Purple: LS4828, green: ATCC29212, orange: JH2-2, and blue: D32.

**Supplemental figure 5. Mutations identified in the c-di-AMP synthesis and degradation genes.** Domain organization (aa) of the **A)** adenylate cyclase CdaA, and the phosphodiesterases **B)** GdpP and **C)** PgpH. Squares express the alignment of the mutated regions from the four different genetic backgrounds. The corresponding mutations are highlighted in red asterisks represent premature STOP codons. The mutated amino acids are bolded and highlighted within the alignment. Putative binding regions, TM domains and active domains are highlighted for each protein. Purple: LS4828, green: ATCC29212, orange: JH2-2, and blue: D32. Line on the top of GdpP protein (B) represents the genetic position (bp) of the gene with a mutation that caused a frameshift.

